# The control of acidity in tumor cells: a biophysical model

**DOI:** 10.1101/2020.03.22.002113

**Authors:** Nicola Piasentin, Edoardo Milotti, Roberto Chignola

## Abstract

Acidosis of the tumor microenvironment leads to cancer invasion, progression and resistance to therapies. We present a biophysical model that describes how tumor cells regulate intracellular and extracellular acidity while they grow in a microenvironment characterized by increasing acidity and hypoxia. The model takes into account the dynamic interplay between glucose and O_2_ consumption with lactate and CO_2_ production and connects these processes to H^+^ and 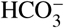 fluxes inside and outside cells. We have validated the model with independent experimental data and used it to investigate how and to which extent tumor cells can survive in adverse micro-environments characterized by acidity and hypoxia. The simulations show a dominance of the H^+^ exchanges in well-oxygenated regions, and of 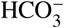 exchanges in the inner hypoxic regions where tumor cells are known to acquire malignant phenotypes. The model also includes the activity of the enzyme Carbonic Anhydrase 9 (CA9), a known marker of tumor aggressiveness, and the simulations demonstrate that CA9 acts as a nonlinear pH_*i*_ equalizer at any O_2_ level in cells that grow in acidic extracellular environments.

**SIGNIFICANCE:** The activity of cancer cells in solid tumors affects the surrounding environment in many ways, and an elevated acidity is a common feature of the tumor microenvironment. In this paper we propose a model of intracellular/extracellular acidity that is linked to cellular metabolism and includes all the main molecular players. The model is reliable, robust and validated with experimental data and can be used as an essential building block of more comprehensive *in silico* research on solid tumors.

## INTRODUCTION

Acid homeostasis in animal tissues is achieved by active dynamic processes. In physiological conditions, the pH of tissues is maintained between 7.35 and 7.45 in spite of constant metabolic acid production by cells. At the microscopic level, cells must finely regulate their own internal pH to around 7.2 to avoid death (1–3). Cellular acid homeostasis is carried out by active transport of acid/base equivalents across the cell membranes into the extracellular spaces.

Dysregulation of pH is a well-known hallmark of solid tumors (1–3). The tissue of solid tumors is characterized by the presence of an irregular network of blood vessels, causing a spatially heterogeneous delivery of nutrients such as glucose and oxygen to tumor cells (1–4). As the consequence, the inner regions of solid cancers that are distant from blood vessels become hypoxic and acidic. Cancer cells adapt to such adverse environments through a series of molecular changes that involve an increased expression of nutrient and ion transporters and enzymes (reviewed in (1, 3, 5)). For example, hypoxia activates the Hypoxia Inducible Factor-1*α* (HIF-1*α*) that up-regulates the transcription of glucose transporters and of enzymes involved in glucose metabolism. Because of hypoxia, glucose is converted mainly to lactic acid through the glycolytic pathway to produce energy under the form of ATP, and the increased production of lactate reduces the pH of the extracellular spaces. A drop in intracellular pH, in turn, increases the activity of lactate and of various ion transporters that collectively contribute to recover intracellular acid homeostasis (1, 3, 5). Hypoxia also causes the increased expression of some membrane-bound enzymes such as Carbonic Anhydrase (CA) that, on the cell surface, catalyzes the hydration of carbon dioxide (CO_2_) to protons (H^+^) and bicarbonate 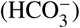 ions. While the H^+^ ions contribute to the acidity of the extracellular milieu, 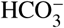 ions can be transported back into the cells and increase the buffering potential of the intracellular environment (1, 3, 5), further contributing to maintain the intracellular pH at normal values.

It has recently been pointed out (1, 3) that changes in the control of intracellular and extracellular acidity in the tissue of solid tumors are associated with many phenotypic changes of cancer cells with important implications in tumorigenesis, cancer progression, cancer diffusion, escape from immune surveillance and resistance to therapies. For example, microscopic examination of the tumor/normal tissue interface shows that peritumoral acidity drives tumor invasion in the surrounding normal tissue, with the regions of highest tumor invasion corresponding to those of lowest pH. In these regions the environmental pH reaches values that are toxic for normal but not for tumor cells (2).

Biophysical models can help to disentangle the intricate relationships between regulatory biochemical networks and give support to the interpretation of experimental evidence which is rapidly accumulating in this field. In this paper we describe a comprehensive biophysical model of the control of acidity in tumor cells. We study the action of key molecular actors in acid homeostasis of cancer cells, and investigate to which extent hypoxia and environmental acidosis influence their behavior. We focus on the dynamic interplay between lactate, proton, bicarbonate transporters and CA enzyme, and their regulation by oxygen and both extracellular and intracellular pH. The model includes the bicarbonate buffer that acts both in the extracellular and intracellular milieux and it incorporates results from our previous modeling efforts concerning tumor cell metabolism (6–8). In particular, our previous models provide values for the rates of glucose and oxygen uptake, lactate and CO_2_ production and lactate/H^+^ transport across cell membranes through specific transporters that have already been validated with experimental data. Finally, we fix the model parameters by combining information from a number of experiments carried out with different tumor cell systems.

## METHODS

### Preliminary considerations and model assumptions

We start from the rather detailed model of tumor cell metabolism and growth that we developed in our previous research (6–8) which successfully reproduces the observed behavior of tumor cells in both liquid (e.g. blood tumors) and solid tumors. In particular, for the current work we have excerpted from that model the part that describes the rates of glucose conversion to lactic acid and oxygen consumption. We remark that the model in (6–8) has been set up with the minimal set of chemical and biochemical pathways that drive the dynamics of metabolism and that are common to most, if not all, tumor cells.

Unlike the metabolic model in (6–8), here we must follow the dynamics of CO_2_, 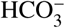 and H^+^, both inside and outside a tumor cell. The inputs of the model are the rates of lactate and CO_2_ production (fig.1) that depend on how cells take up nutrients, such as glucose, and convert them to ATP through the glycolytic and the oxidative phosphorylation pathways. Lactic acid dissociates immediately to lactate and H^+^ ions, and both ions are transported through the cell membrane by means of the bi-directional monocarboxylate transporters MCT (6–8). We remark that this part of the model impacts the rate of change of both intracellular and extracellular pH (from now on pH_*i*_ and pH_*e*_, respectively), and oxygen is assumed to diffuse freely through the cell membrane and its consumption rate is used to determine the rate of CO_2_ production.

**Figure 1:**
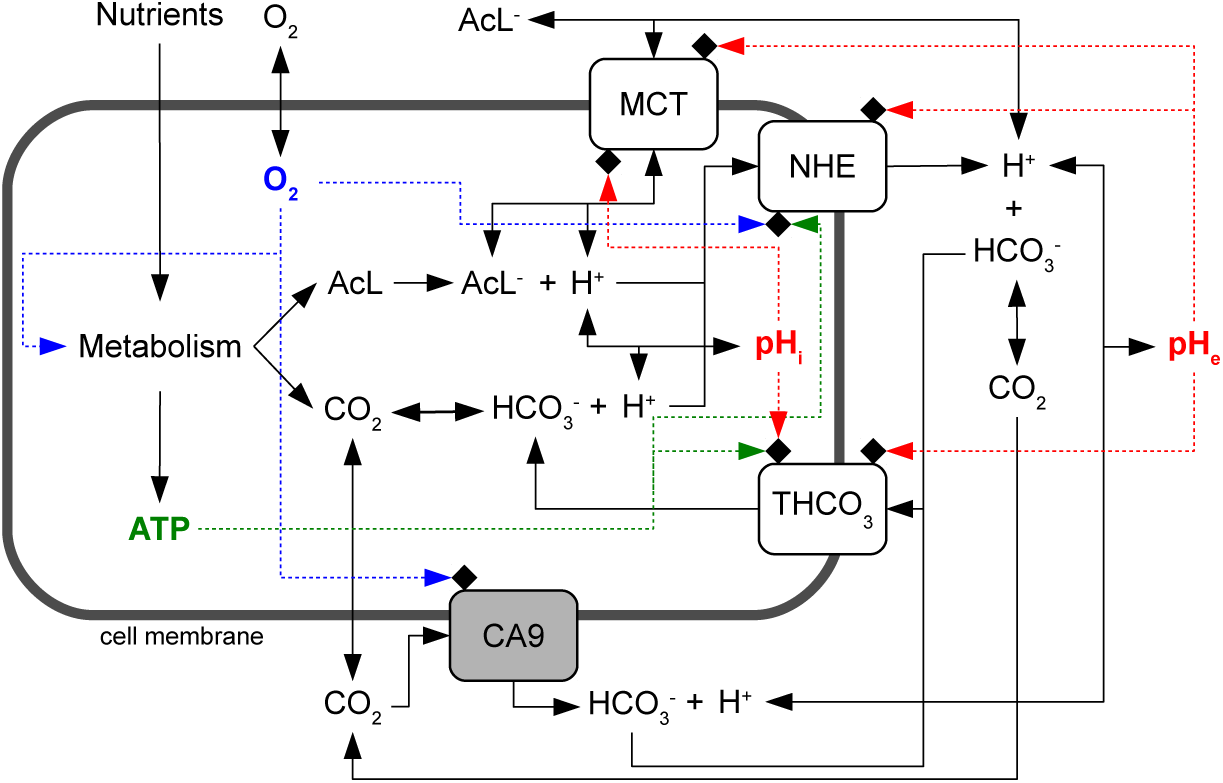
Layout of the model of acidity control in tumor cells. A cell takes up from the environment nutrients and oxygen which are then converted by cell metabolism to lactic acid, CO_2_ and ATP. Lactic acid dissociates to lactate and H^+^ ions, whereas CO_2_ reversibly hydrates to 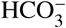 and H^+^. These chemical species diffuse through cell membranes (CO_2_) or are actively transported outside and eventually inside the cell by means of specific protein transporters. We consider monocarboxylate (MCT), sodium-hydrogen exchanger (NHE) and generic bicarbonate (THCO_3_) transporters. We also model the activity of the membrane-bound enzyme Carbonic Anhydrase 9 (CA9). Chemical reactions are indicated by solid lines and the regulatory pathways by dashed lines. Proton concentrations inside and outside the cell are used to compute the intracellular (pH_*i*_) and the extracellular (pH_*e*_) pH. Detailed information on each pathway is given in the main text.

Intracellular H^+^ ions are transported outside the cell by means of unidirectional sodium-hydrogen exchangers NHE (1). Different 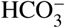 transporters on the other hand are known to drive the flux of bicarbonate ions through the cell membrane. Some of them import or export 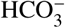 by exchanging Cl^−^ anions and the transport may depend or not on the presence of Na^+^ cations (1). Experimental works, however, have shown that the efficiency of 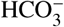 transport in different cell systems is quite similar, and that the import of 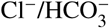 is fundamental in tumor cells where it is dominated by the activity of the Na^+^-dependent 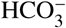 exchanger (9, 10). Therefore, we consider the import activity of a generic 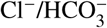 transporter (THCO_3_ in fig.1) which, as a first approximation, assumes the average biochemical characteristics of the Na^+^-dependent 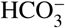 exchanger. We finally model the activity of the membrane-bound Carbonic Anhydrase 9 (CA9) enzyme that catalyzes, on the cell surface, the hydration of CO_2_. This is an important path since CA9 has been found to be expressed by many solid tumors of different histotypes, and its activity has been correlated to tumor progression and growth (11–13).

We model the kinetics of ion transporters, and of CA9 activity as well, with the Michaelis-Menten/Hill formalism that is described by the following general equation:

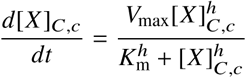

where [*X*]_*C,c*_ is the molar concentration of a given chemical species inside ([*X*]_*C*_) or outside ([*X*]_*c*_) the cell, *V*_*max*_ and *K*_*m*_ are the Michaelis-Menten parameters and *h* is the Hill exponent (*h* > 0).

We assume that:

- CO_2_ can freely diffuse through the cell membrane;
- CO_2_ diffusion is driven by the concentration gradient across the membrane and its only important component is the one directed normally with respect to the cell membrane;
- the diffusion kinetics of charged ions through the cell membrane are much slower than the kinetics of the other processes in which they are involved, and thus the diffusion of charged ions is negligible;
- the mixing of all chemical species in the cell and in the external environment is instantaneous;
- within the short characteristic times of the considered chemical reactions the cell volume is constant.

In this work the variables take the following units for length, mass and time, respectively: *µ*m, pg and s. Molar concentrations (M) have always been converted to mass units by taking into account the volume of the cell (V_*C*_, cell volume is computed by approximating a cell to a sphere of given radius r_*C*_) or of the environment (V_*c*_) and the molecular mass (MW) of chemical species.

### Bicarbonate buffer and initial conditions

Central to the whole scheme of reactions shown in fig.1 is the hydration of CO_2_. It is well known that at physiologic temperature (i.e. ∼ 37°C) carbonic acid dissociates very quickly and represents less than 0.5% of the total carbon dioxide and bicarbonate ion (14). Thus, the hydration of CO_2_ can be approximated by the following chemical reaction:

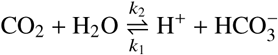

The values of the two rate constants *k*_1_ and *k*_2_ have been determined in cells under standard culture conditions in two independent experiments with good agreement (11, 15). We take the values in (15): *k*_1_ ≃ 0.144 s^−1^ and *k*_2_ ≃ 1.9 · 10^5^ M^−1^ s^−1^.

We compare model outputs with experimental data obtained with cell cultures *in vitro*, in a standard atmosphere at 37°C and 5% CO_2_ at 1 atm pressure. To compute the initial density of CO_2_ dissolved in water under these conditions we use Henry’s law *c* = *k* (*T*)*P* where *c* is the molar concentration of the gas in water, *P* the pressure and *k* (*T*) is a function of temperature

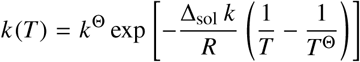

with *T*^Θ^ = 298.15 K, *k*^Θ^ = 3.3 · 10^−4^mol m^−3^ Pa^−1^ and 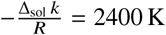 K (see ref. (16) for further details); we find that the initial density of CO_2_ in cell medium under standard culture conditions is:

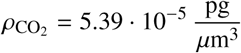

Finally, given the CO_2_ concentration we find the density of 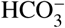 ions from the Henderson-Hasselbach equation:

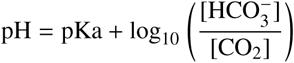

where pKa = − log_10_ (k_1_/k_2_) ≃ 6.12.

Where not otherwise specified, we fixed the standard intracellular and extracellular pH at 7.4, which determines the initial value of the molar concentration of H^+^ ions inside and outside the cells.

### CO_2_ diffusion through the cell membrane

Given the assumptions above, the component of CO_2_ normal to the cell membrane is described by the Fick’s first law:

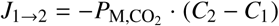

where *J*_1→2_ is the flux from 1 to 2 in units of concentration over time and surface area 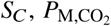 is the permeability of the carbon dioxide and *C*_*i*_ is the concentration of CO_2_ in the *i*-th volume. Since we model cells grown in an incubator at constant CO_2_ pressure, the CO_2_ concentration can reach values far from equilibrium only inside cells because of the oxygen consumption by cell metabolism and of the equivalent CO_2_ production. This means that in the present model there is only a net outward flux of carbon dioxide from cells to the environment. Thus, the net flux of CO_2_ due to diffusion is:

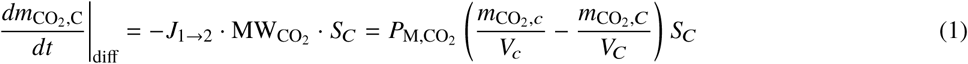

### MCT transporters

The MCTs are a family of bidirectional H^+^ and lactate co-transporters expressed at the cell membrane and their activity has been shown to depend on the pH values on both sides of the cell membrane (see refs. (6–8) and references therein). We model their activity with parameter values extrapolated from experimental observations (6–8) and we use the following equations and parameters to describe the rate of transport of H^+^ inside and outside the cell:

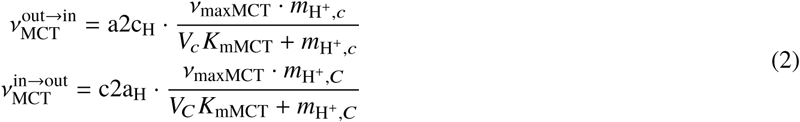

where 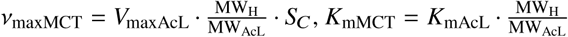 and where the ratio of molecular weights is used to rescale the equations from concentrations to masses.

In eqns.2, a2c_H_ and c2a_H_ depend, respectively, on extracellular and intracellular pH, and phenomenologically describe the dependency of MCT transport activity on acidity (for a complete analysis see (6–8)):

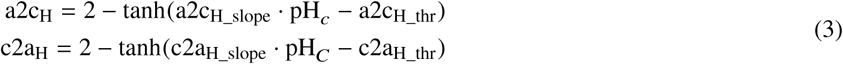

### NHE transporters

Sodium-hydrogen exchangers (NHE) are membrane transport proteins that exploit the influx of Na^+^ to export H^+^ ions. The sodium concentration gradient is maintained by the ATP-dependent Na^+^/K^+^ pump (17, 18) so that the activity of NHE indirectly depends on ATP availability. This implies that as long as ATP is available the flux of H^+^ due to NHE is essentially unidirectional. It has also been reported that NHE activity is inhibited by hypoxia (10, 17) and that, in the long-term, hypoxia inhibits the expression of NHE proteins. Energy and oxygen tune NHE activity and as in the previous model of tumor cell metabolism and growth (6–8), here we take into account these regulatory circuits by means of the two variables SensATP and SensO_2_ that assume real values in the interval [0, 1].

Experimental observations indicate that NHE activity is described by a Hill equation (9, 19, 20) and hence the unidirectional flux of H^+^ from the cell to the environment due to NHE transport is modeled by the equation:

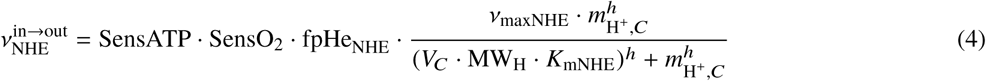

where *ν*_maxNHE_ = V_maxNHE_ · *S*_*C*_ and fPHe_NHE_ is a phenomenological function that tunes the activity of NHE transport as a function of extracellular pH:

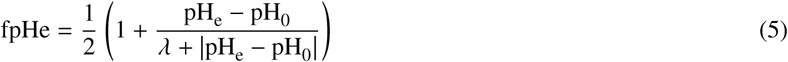

Indeed, it has been observed that extracellular acidity enhances H^+^ transport through NHE (9, 17, 21). In the Supporting Material we discuss how we fix parameter values and define the function fPHe on the basis of experimental observations.

### Transport of bicarbonate ions

As discussed above, we model the activity of a generic bicarbonate ion importer (THCO3). The Na^+^-dependent 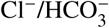 exchanger appears to dominate 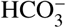 fluxes in tumor cells (9, 10), and therefore we take this transporter as a reference to set the values of parameters and fix general biochemical characteristics. This is an important part of the model, because it has been shown that tumor cells do actively import 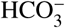 ions to buffer their internal pH (9, 10), and that this is a common property of different cancer cells. Experimental studies have demonstrated that 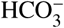 import is regulated by both intracellular and extracellular pH but not by hypoxia and that the transport follows a simple Michaelis-Menten kinetics. In the scientific literature there are no indications, as far as we can tell, that 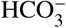 transport depends on ATP availability. However, just as observed for proton export by NHE transporters, 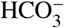 transport proceeds by parallel fluxes of ions, like Na^+^ and Cl^−^, along their electrochemical gradients that are actively maintained by cells through energy-consuming paths. Thus, it is likely that even 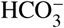 transport is controlled by ATP availability, albeit indirectly. On the basis of these considerations we model 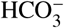 import as follows:

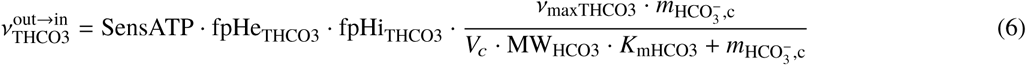

where where *ν*_maxTHCO3_ = V_maxTHCO3_ · *S*_*C*_ and the two functions fpHe_THCO3_ and fpHi_THCO3_ phenomenologically describe how 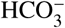 import is affected by extracellular and intracellular pH, respectively. These functions have been fit to actual experimental data (see the Supporting Material) and are modeled by the following equations:

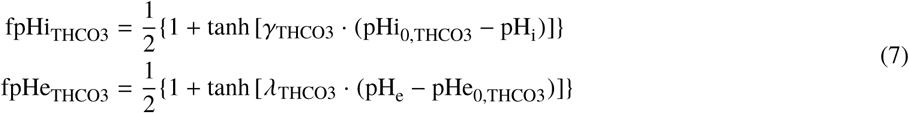

### Activity of Carbonic Anhydrase 9

The enzyme CA9 is expressed by cells of many different solid tumors, and in general its expression correlates with cancer aggressiveness and poor therapeutic outcome (11–13). It is a membrane-tethered enzyme and it is mainly found at the external surface of cells where it catalyses the hydration of CO_2_ (11–13). Importantly, its expression is regulated by hypoxia and indeed CA9 is a marker of hypoxia (22). Again, experimental observations show that CA9 activity follows a Michaelis-Menten kinetics.

Thus:

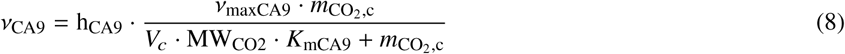

where where *ν*_maxCA9_ = V_maxCA9_ · *S*_*C*_ and h_CA9_ is a phenomenological functions that describe how hypoxia tunes CA9 expression:

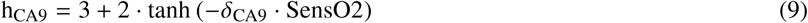

This is a function of the fraction of available oxygen which, in our model, is defined by SensO2, and it describes the fold change in CA9 expression as observed in actual experiments (see the Supporting Material).

### The full model and its numerical integration

The full model is represented by the following set of differential equations:

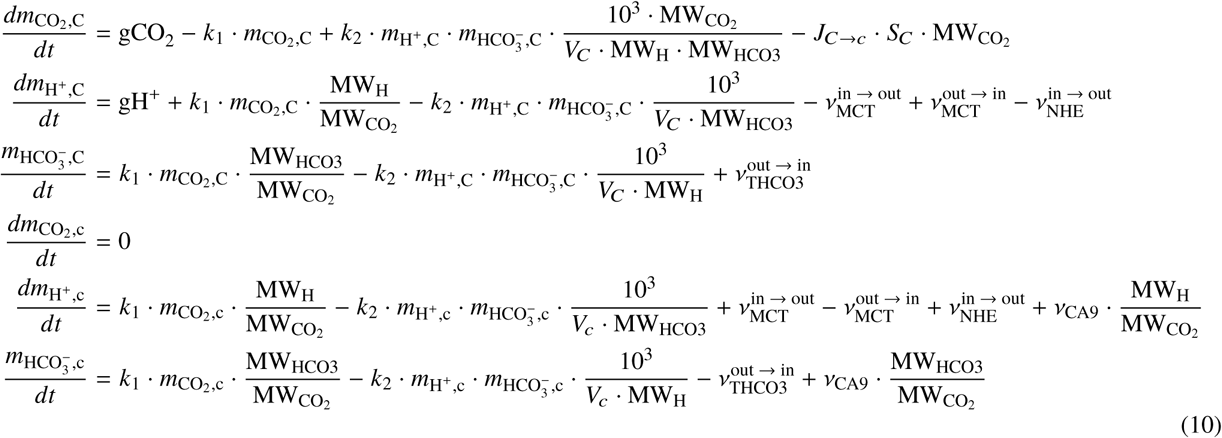

where gH^+^=gAcL·MW_H_/MW_AcL_ and 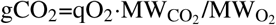 are, respectively, the rates of H^+^ and CO_2_ production that are proportional to the rate of lactate production gAcL and oxygen consumption qO_2_ as defined in our previous work (6–8), and all the other rates, and regulatory functions, are given in equations 1–9. The multiplicative factor 10^3^ that appears in the right-hand side of equations 10 above comes from the conversion of standard molar concentration units to the units used here where masses are expressed in pg and volumes in *µ*m^3^.

The system of differential equations 10 is nonlinear and stiff because it incorporates processes with different kinetics, from the fast kinetics of CO_2_ hydration and diffusion to the relatively slow kinetics of ion transport and enzyme activity. The system cannot be solved analytically and appropriate numerical approaches are required. We previously investigated this aspect within the context of complex large-scale biophysical models (23) and found that the implicit Euler method is well-suited for the numerical integration of models of this kind. We solved the discretized system of differential equation 10 using the implicit Euler algorithm followed by the Newton-Raphson method to solve numerically the resulting system of nonlinear equations. The code has been implemented in C++ using the computational framework provided by the GNU Scientific Library (24). We used the standard Newton-Raphson solver *gsl_multiroot_fsolver_dnewton* and the *gsl_multiroot_test_residual* library to test the convergence of the algorithm (threshold *ϵ* < 10^−6^) within a maximum number of iterations fixed at *N*_max_ = 1000.

## RESULTS

The model defined by the set of differential equations 10 has several parameters. We extensively searched the scientific literature to find their values, and when these values were not directly available they were obtained by fit of specific equations to reported experimental data. Experimental evidence was also used to model regulatory functions given by equations 3, 5, 7 and 9 that tune the activity of transporters and CA9 enzyme as the function of local pH, ATP and/or oxygen availability. The full strategy is detailed in the Supporting Material and all parameter values are listed in Table 1.

**Table 1:**
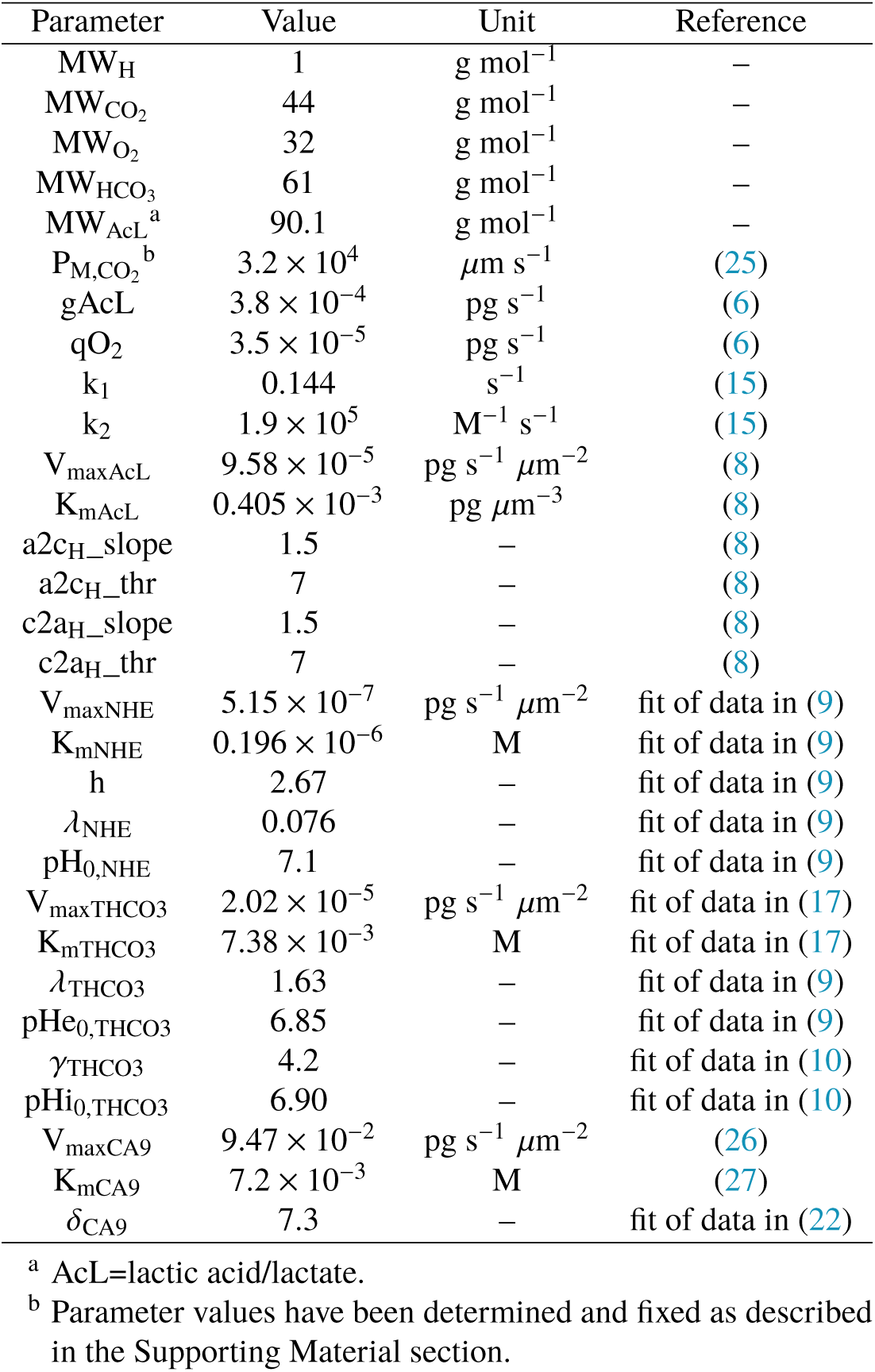
Values of model parameters

| Parameter | Value | Unit | Reference |
| --- | --- | --- | --- |
| $MW_H$ | 1 | $\text{g mol}^{-1}$ | – |
| $MW_{CO_2}$ | 44 | $\text{g mol}^{-1}$ | – |
| $MW_{O_2}$ | 32 | $\text{g mol}^{-1}$ | – |
| $MW_{HCO_3}$ | 61 | $\text{g mol}^{-1}$ | – |
| $MW_{AcL}^a$ | 90.1 | $\text{g mol}^{-1}$ | – |
| $P_{M,CO_2}^b$ | $3.2 \times 10^4$ | $\mu\text{m s}^{-1}$ | (25) |
| $g_{AcL}$ | $3.8 \times 10^{-4}$ | $\text{pg s}^{-1}$ | (6) |
| $q_{O_2}$ | $3.5 \times 10^{-5}$ | $\text{pg s}^{-1}$ | (6) |
| $k_1$ | 0.144 | $\text{s}^{-1}$ | (15) |
| $k_2$ | $1.9 \times 10^5$ | $\text{M}^{-1} \text{s}^{-1}$ | (15) |
| $V_{\max AcL}$ | $9.58 \times 10^{-5}$ | $\text{pg s}^{-1} \mu\text{m}^{-2}$ | (8) |
| $K_{mAcL}$ | $0.405 \times 10^{-3}$ | $\text{pg } \mu\text{m}^{-3}$ | (8) |
| $a_{2c_H\_slope}$ | 1.5 | – | (8) |
| $a_{2c_H\_thr}$ | 7 | – | (8) |
| $c_{2a_H\_slope}$ | 1.5 | – | (8) |
| $c_{2a_H\_thr}$ | 7 | – | (8) |
| $V_{\max NHE}$ | $5.15 \times 10^{-7}$ | $\text{pg s}^{-1} \mu\text{m}^{-2}$ | fit of data in (9) |
| $K_{mNHE}$ | $0.196 \times 10^{-6}$ | M | fit of data in (9) |
| $h$ | 2.67 | – | fit of data in (9) |
| $\lambda_{NHE}$ | 0.076 | – | fit of data in (9) |
| $pH_{0,NHE}$ | 7.1 | – | fit of data in (9) |
| $V_{\max THCO_3}$ | $2.02 \times 10^{-5}$ | $\text{pg s}^{-1} \mu\text{m}^{-2}$ | fit of data in (17) |
| $K_{mTHCO_3}$ | $7.38 \times 10^{-3}$ | M | fit of data in (17) |
| $\lambda_{THCO_3}$ | 1.63 | – | fit of data in (9) |
| $pHe_{0,THCO_3}$ | 6.85 | – | fit of data in (9) |
| $\gamma_{THCO_3}$ | 4.2 | – | fit of data in (10) |
| $pHi_{0,THCO_3}$ | 6.90 | – | fit of data in (10) |
| $V_{\max CA9}$ | $9.47 \times 10^{-2}$ | $\text{pg s}^{-1} \mu\text{m}^{-2}$ | (26) |
| $K_{mCA9}$ | $7.2 \times 10^{-3}$ | M | (27) |
| $\delta_{CA9}$ | 7.3 | – | fit of data in (22) |

<sup>a</sup> AcL=lactic acid/lactate.
<sup>b</sup> Parameter values have been determined and fixed as described in the Supporting Material section.

Once determined, parameter values were fixed and no further tuned to adapt model outputs to data. This means that the model has no free parameters and is strictly predictive. As explained in the next section, for validation purposes we first used it to predict how the intracellular pH (pH_*i*_) varies when cells are grown into environments with increasing acidity.

### Model validation with independent experimental data

Model validation was performed with independent experimental data, i.e. data that were not used to set parameter values. To this end we used the data in the paper by Song et al. (28). In this paper Song et al. investigated the dependence of pH_*i*_ on pH_*e*_ in SCK cells (human choloangiocarcinoma cell line) in standard *in vitro* cultures. Unfortunately, the radius of SCK cells is not reported nor, to the best of our knowledge, it has been measured previously. This is important because our model equations take into account both cell volume (see eqns.1–10) and the cell surface (see e.g. CO_2_ diffusion, eq.1) that are both computed from cell radius under the assumption that cell geometry can be approximated by a sphere.

Figure 2 shows the model prediction for intracellular pH vs. cell size, under standard culture conditions. At equilibrium there is a difference of ≈ 0.1 in pH between small and large cells (r_*C*_ = 5.5 and 8.0 *µ*m, respectively, i.e. a volume ratio of ≃ 3) but pH_*i*_ levels reach values that have actually been observed in tumor cells (28). With the initial conditions discussed above, the simulations approach equilibrium quite fast and this indicates that the numerical solution of model equations is stable.

**Figure 2:**
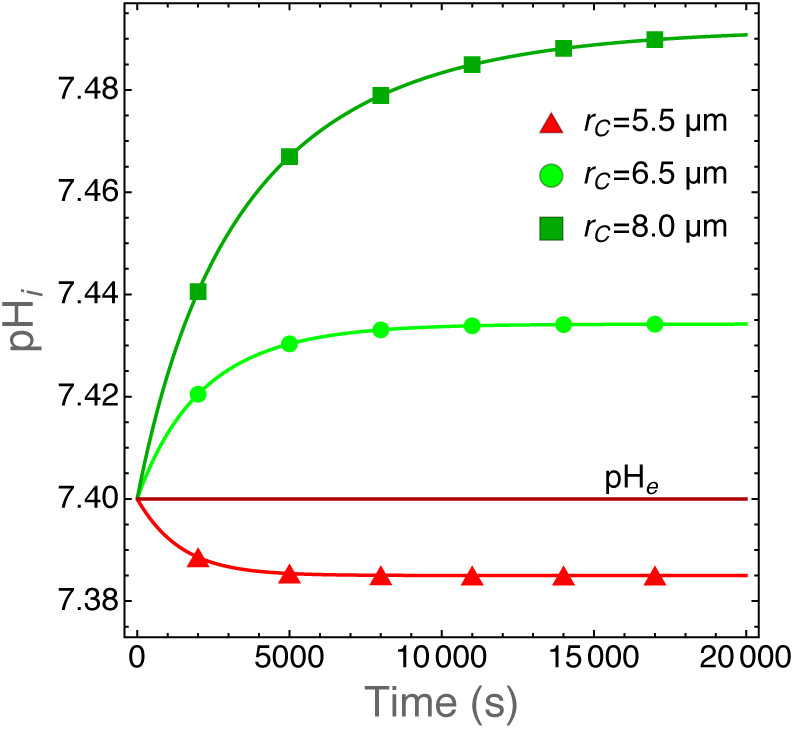
Plot of pH_*i*_ as the function of time for cells with the indicated cell radii. We take r_*C*_ values that are in the observed range for human tumor cells (29). The model equations have been solved with the parameter values listed in Table 1. After an initial transient, pH_*i*_ reach an equilibrium at physiological values and this shows that the model (and its numerical solution) is stable and provides quantitative results in good agreement with actual experimental observations. We also plot pH_*e*_ for comparison. The extracellular pH does not vary because these runs were carried out for a limited time span and for cells growing in a large volume (1 mL) filled with fresh medium at physiological pH to mimic standard experimental conditions.

The model predictions for pH_*i*_ values in SCK cells grown in media with increasing acidity are shown in figure 3. We ran simulations with varying cell radius within a range of values which is reasonable for tumor cells, i.e. between 4.5 and 9 *µ*m (29) and computed pH_*i*_ at equilibrium. As shown in figure 2 the numerical solutions approach equilibrium with slower kinetics for increasing cell radii. We chose a conservative criterion to define the equilibrium condition and we halted the simulations when ΔpH_*i*_/Δ*t* < 10^−5^ was reached. In these simulations, the volume of the environment was set to V_*c*_ = 10^12^ *µ*m^3^ = 1mL, i.e. large enough to assure nearly constant pH_*e*_ values throughout the simulation runs. Figure 3 shows that model predictions are in excellent agreement with the experimental data.

**Figure 3:**
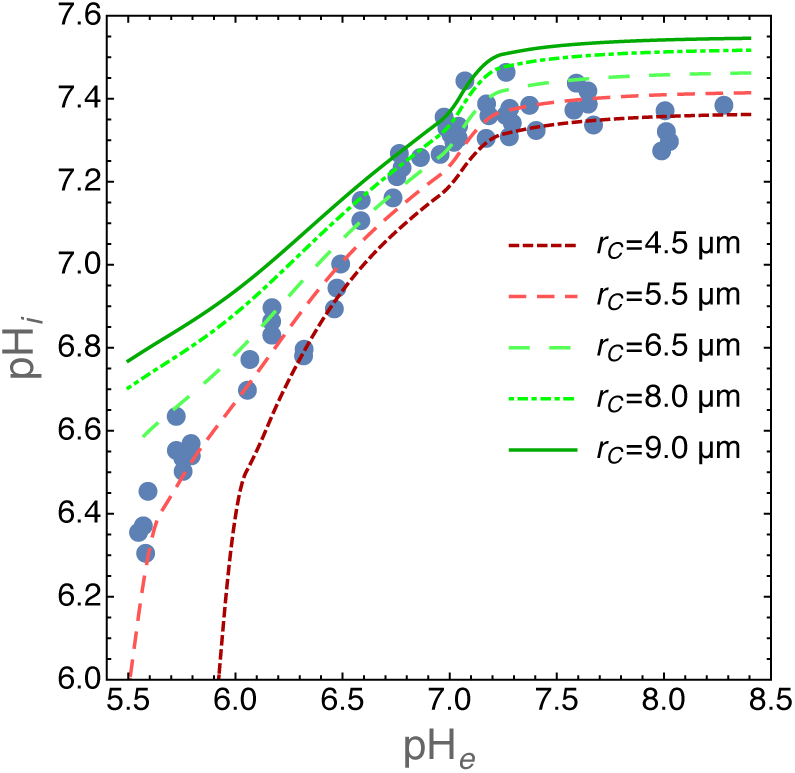
Plot of pH_*i*_ for SCK cells grown in media with different pH_*e*_ values. Experimental data have been redrawn from figure 2 in (28) (closed circles). The lines show pH_*i*_ values at equilibrium as predicted by our model for the indicated cell radii. It is important to note that these are not fits because our model does not have free parameters. Equilibrium was reached at ΔpH_*i*_/Δ*t* < 10^−5^. The volume of the environment was set at V_*c*_ = 10^12^ *µ*m^3^ = 1mL.

### Contribution of NHE and THCO3 transporters to pH_*i*_ in normoxic or hypoxic environments

We have used the model to study the biochemical mechanisms that allow tumor cells to survive to adverse environments. We have investigated the role of NHE and THCO3 transporters in the control of intracellular acidity by tumor cells exposed to normoxic or hypoxic environments. We ran several simulations by alternatively switching off the activity of NHE and THCO3 transporters, i.e. by setting the respective *ν*_max_ parameters to 0. The results are shown in figure 4 where we plot the pH_*i*_ values at equilibrium (see the previous section) as the function of environmental pH for cells grown under standard oxygen level or at 0.1 fraction thereof.

**Figure 4:**
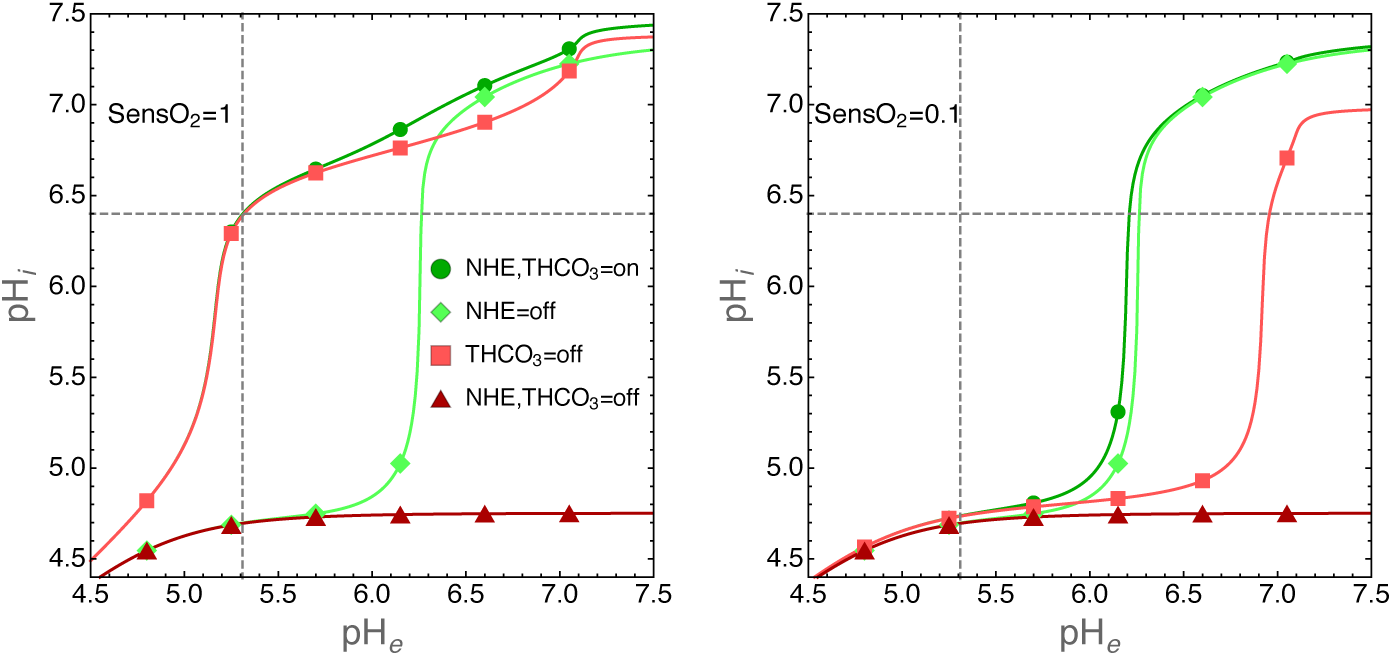
Contribution of NHE and THCO3 transporters to pH_*i*_ in normoxic (left panel) or hypoxic (right panel) environments. Simulations were run with the following parameters: cells radius r_*C*_ = 6.5*µ*m and environmental volume V_*c*_ = 10^12^ *µ*m^3^. The intracellular pH was calculated at equilibrium (see also the legend to figure 3) as the function of the indicated pH_*e*_ values. The activity of NHE and THCO3 transporters was switched off by setting the respective *ν*_max_ parameters to 0. Environmental oxygen levels were tuned by setting the SensO2 parameter to 1 or to 0.1 (see the Methods section and the Supplementary Material for details). In both panels, dashed lines have been drawn to show the pH_*e*_ value at which pH_*i*_ = 6.4, a value largely compatible with cell life (see also the experimental data in figure 3 for a comparison).

The simulations clearly show that under normoxic condition the contribution of the THCO3 transporter to pH_*i*_ is negligible. Under this condition pH_*i*_ is maintained to physiological levels thanks to the activity of NHE transporter that export H^+^ ions outside the cells. On the contrary, THCO3 activity dominates in hypoxic environments.

### Role of Carbonic Anhydrase 9

As previously noted by Swietach and colleagues (11) pH_*i*_ regulation is not affected by CA9 expression in isolated tumor cells, but its role becomes important when cells are grown as three-dimensional aggregates (tumor spheroids). When expressed by cells grown as tumor spheroids CA9 induces a near uniform intracellular pH throughout the structure (11), an observation that was explained by diffusion-reaction modeling as follows: CA9 coordinates pH_*i*_ spatially by facilitating CO_2_ diffusion in the unstirred extracellular space of the spheroid (11). This intriguing conclusion, supported by experimental evidence, suggests that CA9 activity becomes important for the control of pH_*i*_ by tumor cells at critical sizes of the extracellular volume. We tested this hypothesis with our model, and the results are shown in figure 5.

**Figure 5:**
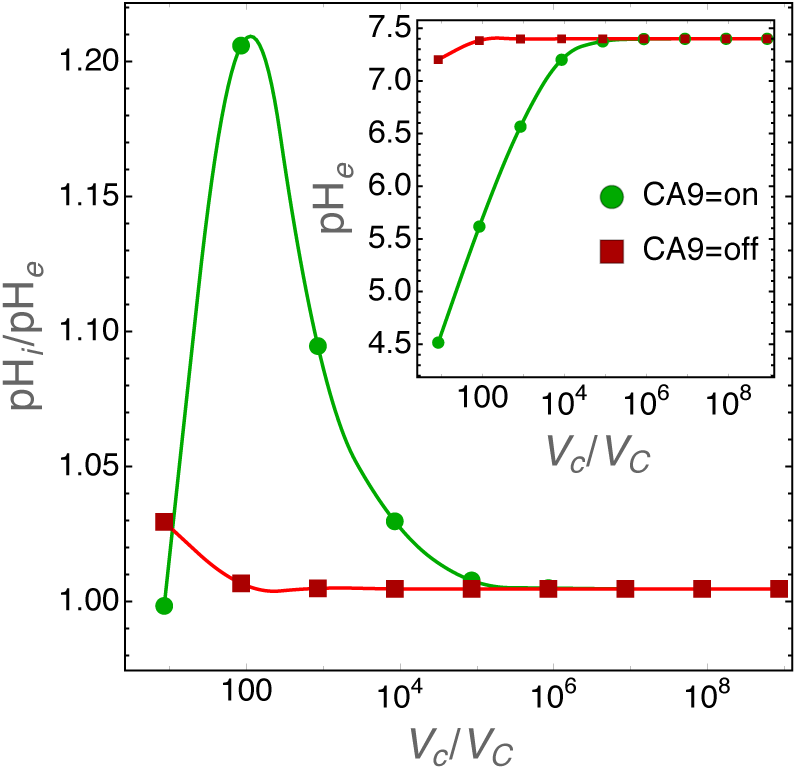
pH_*i*_ regulation by CA9 for decreasing size of the extracellular volume. Cell radius was set to the average size of 6.5 *µ*m. The inset shows pH_*e*_ values and the main panel pH_*i*_ to pH_*e*_ ratio for varying V_*c*_/V_*C*_ values (i.e. ratio of extracellular to cell volumes) when CA9 activity is turned on or off. In these simulations the extracellular environment is physically closed, i.e. the extracellular volume is unstirred and the diffusion of chemical species toward an “external reservoir” is not allowed.

The role of CA9 in pH_*i*_ regulation starts to become important at the extracellular to cell volume ratio V_*c*_/V_*C*_ ≈10^4^ and reaches a maximum at V_*c*_/V_*C*_ ≈ 100. It is important to note that we simulated cells that grow in a closed environment. This means that at small extracellular volumes the acidity of the environment becomes too high and pH_*i*_ runs out of control (see also figure 4). However, the results in figure 5 show that when V_*c*_/V_*C*_ ≈100 and CA9 is active the extracellular pH at equilibrium is around 5.5 and pH_*i*_ ≈6.6, well within the physiological range.

Simulations in figure 5 do not take into account the oxygen levels in the tumor environment. As discussed above (see the Methods section) CA9 expression is regulated by hypoxia (22) and thus it is interesting to investigate how pH_*i*_ is regulated by cells growing in small environments, i.e. when the CA9 role is not negligible, and when O_2_ levels are lower and lower. Figure 6 shows that when pH_*e*_ ≥ 5.8, CA9 acts as a nonlinear pH_*i*_ equalizer at any O_2_ levels.

**Figure 6:**
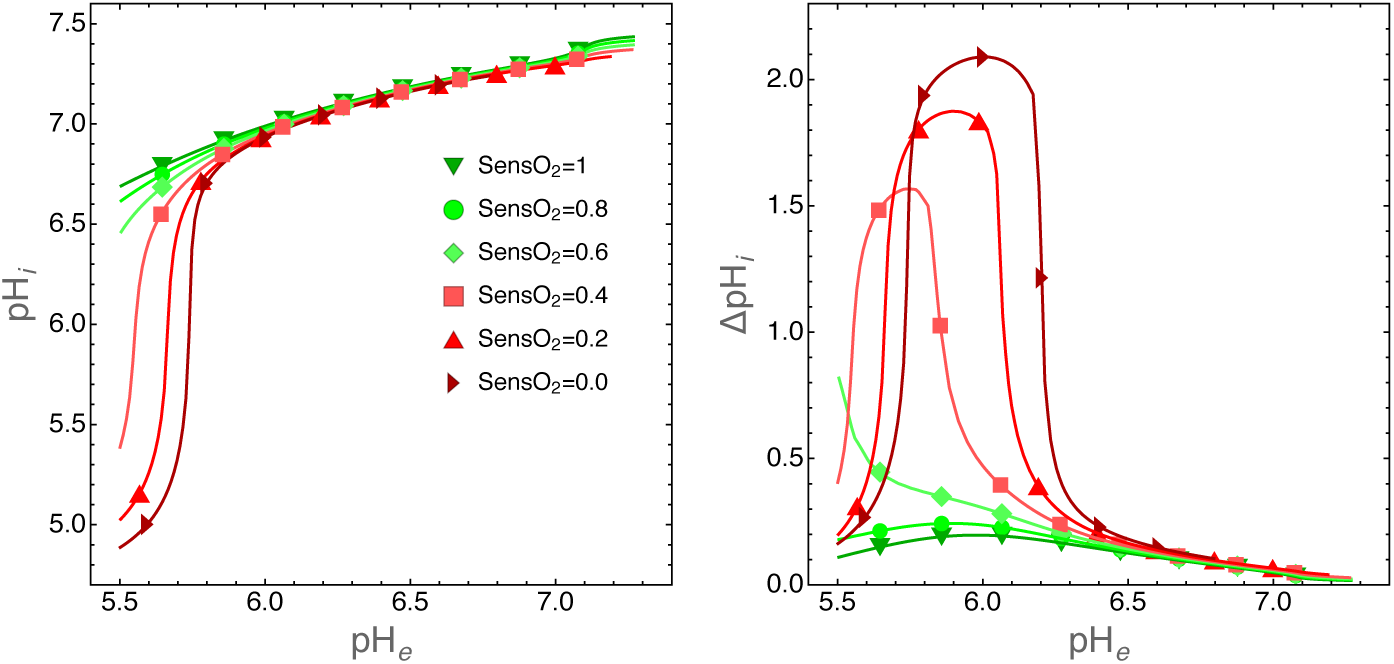
Role of CA9 on pH_*i*_ regulation for cells grown in a small environment with decreasing oxygen levels. In these simulations the extracellular volume was set to V_*c*_ = 10^5^ *µ*m^3^ and cell radius to r_*C*_ = 6.5*µ*m so that V_*c*_/V_*C*_ ≈80. Left panel: plot of pH_*i*_ at equilibrium as the function of pH_*e*_ for the indicated fractions of environmental O_2_. Right panel: same simulations as those shown in the left panel, but here we plot ΔpH_*i*_ = pH_*i*,CA9=on_ − pH_*i*,CA9=off_, i.e. the difference in pH_*i*_ when CA9 is turned on or off. This plot clearly shows the nonlinear character of CA9 activity in the regulation of pH_*i*_.

## DISCUSSION

We have developed a biophysical model to explore the complex molecular mechanisms that allow tumor cells to regulate both intracellular and extracellular acidity, but we are not alone, other modeling efforts have tried to capture the essential features of the biochemical pathways that lead to acid homeostasis in tumor cells (see e.g. (30–33)). We have taken the remarkable models described in (32) and (33) as our starting point, because of their direct applicability to the analysis of experimental data. The former provides a fully tractable quantitative description of the interplay between H^+^ and 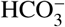 transporters with Na^+^/K^+^-ATPase and Na^+^, K^+^ and Cl^−^ ion fluxes, while the latter investigates the interaction of MCT transporters and CA9. We go a few steps further and model the network of important paths that connect together cell metabolism and hypoxia with transport of H^+^ and 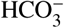 ions and CA9 activity (see figure 1). The coupling of ion transport mechanisms with metabolism and hypoxia is essential if we want to understand how tumor cells grow and shape their microenvironment, an interplay that is of fundamental importance for the adaptation and evolution of cancer cells within a solid tumor.

We remark that with the model described here we are able to give a quantitative assessment of the importance of specific molecular mechanisms. For instance, simulations show that H^+^ efflux from tumor cells dominates the control of intracellular acidity in normoxic environments, whereas 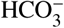 import in hypoxic tumor areas (in our simulation where the fraction of oxygen decrease to 0.1 of standard values). Experiments have shown that in *in vivo* tumor micro-environments oxygen reaches 10% of its normal value at a distance of ≈ 150*µ*m, i.e. ≈ 10 cell diameters, from blood vessels (34). Thus, within this short distance the control of pH_*i*_ is attained by tumor cells through a switch from H^+^ export to 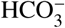 import pathways. This observation gives further support to recent work that has shown that inhibition of 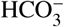 fluxes inhibits the growth of experimental tumors by increasing intracellular acidity and cell death (35). When we recall that the hypoxic regions are those where tumor cells show higher resistance to therapies, such as e.g. radiotherapy, then we see that approaches that aim at inhibiting 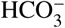 fluxes would target the very cells that colonize the inner tumor regions and that would otherwise be resistant to therapies, and improve cancer control.

Finally, the model singles out the important role of CA9. The simulations show that CA9 acts as a nonlinear pH_*i*_ equalizer at any O_2_ level in cells that grow in acidic extracellular environments. This result is in agreement with the experimental observations by Swietach and colleagues (11), collected with tumor spheroids. They observed near-uniform pH_*i*_ values throughout the spheroid structure due to CA9 activity in spheroids grown up to ≈ 500*µ*m diameter. It has long been recognized that tumor spheroids of this size show steep gradients of oxygen with fractions that go as far down as 0 at the center of the spheroid (36). Our simulations show that this is due to the concerted action of CA9 and of hypoxia that up-regulates CA9 expression. These two mechanisms collectively help cells to keep their intracellular pH under control because of increased 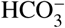 production followed by 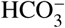 import through THCO3 transporters.

## CONCLUSION

While tumor cell adaptation and survival to extreme microenvironments are key concepts in oncogenesis (1–3), we remark that acid homeostasis is central to cellular adaptation in a much wider context. Active transport of acid/base equivalents across cell membranes into the extracellular spaces may cause transient and rapid changes of microenvironmental and cellular pHs like those observed for other ions involved in cell signalling. Indeed, pH transients have been shown to be important in intra- and inter-cellular communication in the nervous system and are known to affect a number of essential functions, like e.g. neuronal excitability and synaptic transmission (37). This in turn implies that animal cells could sense and adapt to pH changes. The underlying molecular mechanisms are still not well understood, but the role of G-protein coupled receptors in proton sensing is increasingly investigated also in relation to pathological conditions, besides cancer, that result in an increased extracellular acidity, such as infarction and inflammation (38). We conclude that our model can be used as an essential building block of more comprehensive *in silico* research on solid tumors, but it may also help understanding how other cells can sense and dynamically adapt to pH changes.

## AUTHOR CONTRIBUTIONS

RC and EM designed the research. RC found parameter values. NP and EM wrote C++ code. NP and RC carried out simulations. RC and EM wrote the article. All authors critically discussed results and revised the article.

